# Host cohort has a larger impact on the gut microbiomes of mealworms and superworms than does the ingestion of polyethylene or polystyrene

**DOI:** 10.64898/2026.04.25.720593

**Authors:** Sabhjeet Kaur, Jithin S. Sunny, Sydnie A. Stinson, Mia A. Rondinelli, Jeremie Alexander, Aidan Vint, Rebecca T. Doyle, Laura A. Hug, George C. diCenzo

**Author notes:** **Corresponding author:** George C. diCenzo.

## Abstract

Beetles (order Coleoptera) are a diverse insect group whose associated microbial communities remain understudied despite their ecological, commercial, and biotechnological importance. Larvae of the darkling beetle species *Tenebrio molitor* (mealworms) and *Zophobas atratus* (superworms) have gained attention for their reported ability to ingest and possibly degrade plastics such as polyethylene and polystyrene. This degradation is believed to be mediated by enzymes from their gut microbiota, yet the underlying microbial mechanisms remain unclear. Using deep shotgun metagenomics, we generated comprehensive gut metagenomes for mealworms and superworms, recovering 53 and 100 high-quality metagenome-assembled genomes (MAGs), respectively. We found that the gut microbiomes of both insect species are dominated by bacteria from the phyla *Bacillota* and *Pseudomonadota*, with superworms trending towards greater bacterial diversity than mealworms. Comparing gut bacterial communities of insects fed polyethylene or polystyrene versus controls revealed no clear plastic ingestion effects beyond those attributable to starvation. On the other hand, insect cohort contributed substantially to variation in bacterial community composition, explaining 23% and 33% of the total variation in mealworms and superworms, respectively. Finally, we computationally identified numerous secreted proteins with sequence similarity to enzymes previously implicated in polyethylene and polystyrene degradation, and we provide support for dye decolorizing peroxidase (DyP)-type peroxidases as being the enzyme class mostly likely to initiate plastic oxidation in the digestive tracts of mealworms and superworms. Overall, this study provides high-quality metagenomes and MAGs for mealworms and superworms, revealing substantial under-explored microbial biodiversity, and highlighting DyP-type peroxidases as promising targets for future plastic biodegradation studies.

## INTRODUCTION

Beetles (order: Coleoptera) are the largest and most diverse group of insects [1]. Although many studies focus on the negative impact of beetles as pests in forest and agriculture [2, 3], they play pivotal roles in ecology as plant pollinators and scavengers due to their biodiversity and widespread nature [4]. Beetles, both adults and their larvae, are important commercially as feeder insects for pets such as reptiles and fish, and they are among the more than 2,000 species of insects consumed by humans globally as a source of protein [5]. As a group, beetles are thought to host extensive microbial biodiversity, with some researchers estimating that beetles collectively host over 3 million unique bacterial species [6, 7]. Beetle microbiomes may therefore host many biotechnologically relevant microbes of value to the pharmaceutical industry [8], waste management [9], and other industries. However, despite the potentially vast biodiversity and functional potential associated with beetle gut microbiomes, metagenomic studies of these communities remain scarce, leaving these systems comparatively understudied.

The larvae of the darkling beetle (family: Tenebrionidae) species *Tenebrio molitor* and *Zophobas atratus* (syn. *Zophobas morio*), known as mealworms and superworms, respectively, are two examples of industrially reared beetle species that have gained significant scientific attention. In addition to being edible insects and sold as feeder insects for pets, mealworms and superworms are notable due to reports that they will chew, and ingest, various plastic polymers including polyethylene (PE), polystyrene (PS), polypropylene, and polyvinyl chloride (PVC) [10–14]. Several studies have additionally provided evidence that mealworms and superworms not only ingest the plastics but can degrade and digest a portion of it [10, 15–19], although other studies provide contradictory evidence suggesting that only the additives (not the plastic polymers themselves) are oxidized [20]. Mealworms co-fed antibiotics appear unable to digest the plastics [18, 21–23], suggesting that the apparent degradation of plastic polymers by these insects is at least partially dependent on enzymes encoded by bacteria present in their digestive tracts. These findings have led to scientific interest in understanding how these beetles, and their microbiome, may degrade plastics to inform the development of novel biotechnologies for plastic waste management [13, 24].

Several studies have examined how ingestion of plastics shifts the microbial community composition of mealworm and superworm digestive tracts. 16S rRNA gene amplicon sequencing studies have observed an increase in the relative abundance of several genera, including but not limited to *Citrobacter* and *Enterobacter* (phylum *Pseudomonadota*), *Lactococcus* and *Enterococcus* (phylum *Bacillota*), and *Spiroplasma* (phylum *Mycoplasmatota*) [15–19], which could suggest members of these taxa are able to catabolize the ingested plastics. However, it is important to note that prior to our study, only a few published studies (e.g., [25]) include a comparison with unfed (i.e., starved) insects, and thus for most studies, we cannot differentiate whether the observed enrichment is directly due to the plastics or is a result of the insects experiencing starvation. In addition, unlike shotgun metagenomics, 16S rRNA gene amplicon sequencing studies are unable to provide mechanistic insight into the apparent plastic biodegradation by the gut microbiota.

We are aware of only four studies using shotgun metagenomics to characterize the gut microbiome of mealworms or superworms. Sun et al. [26] used metagenomics to characterize how PS exposure impacted the gut microbiome of superworms, although fewer then ∼10 million paired reads were generated per sample and only 11 MAGs were generated. Similarly, Xu and Dong [27] used metagenomics to characterize the impact of PVC on the gut microbiome of mealworms, generating fewer than 10 million bacterial reads per sample without performing binning to obtain MAGs. In a study unrelated to plastic biodegradation, Leierer et al. [28] used metagenomics to characterize the microbiome of mealworm colonies, generating an average of fewer than 10 million paired reads per sample, and with a focus on analysis of the raw reads without assembling metagenomes. Lastly, Klauer et al. used Pacific Biosciences sequencing to generate comprehensive metagenomic data for mealworms with or without PE exposure, obtaining up to 23 high-quality MAGs per sample [29]. However, several of their datasets appear to principally contain host DNA, and the data were used primarily to investigate the abundance of a specific gene family, with no community level analyses reported. Overall, the limitations of prior shotgun metagenomics studies of mealworms and superworms means that much remains to be discovered about the biodiversity and functional potential of the gut microbiomes of these insects and the impacts of ingesting plastics.

To overcome these limitations, we used shotgun metagenomics (∼64 million read pairs per sample) to generate high-quality, reference gut metagenomes for mealworms and superworms, reporting dereplicated sets of 53 and 100 high-quality metagenome-assembled genomes (MAGs) for mealworms and superworms, respectively. In addition, we attempted to investigate how the gut microbiome responds to ingestion of PE or PS, with starved insects used as a control, and computationally screened the metagenomes for oxidative enzymes potentially relevant for early steps in the biodegradation of PE or PS.

## MATERIALS AND METHODS

### Insect colonies and experimental design for metagenomic sequencing

Each week for three weeks, 500 mealworms (*T. molitor*) and 250 superworms (*Z. atratus*) were purchased from a local pet store in Kingston, ON, Canada. Each weekly batch of insects of the same species, hereafter referred to as a cohort, was placed into a single Starfrit 1.6 L LocknLock Glass Container (Starfit catalog number 095010004NEW1) with oatmeal, wheat bran, and carrot slices to acclimatize before putting them on specific diets. A hole (diameter of ∼6.5 cm) was cut in the lid of each container and covered with an aluminum insect screen to allow for air exchange. Insect colonies were maintained at room temperature and ambient moisture for two days, at which point insects were transferred into new Starfrit 1.6 L containers. Four containers were prepared per insect species, with 100 mealworms or 50 superworms added per container; fewer superworms were added per container to account for their larger size. Each of the four containers per species contained one of the following and nothing else: (i) oatmeal and bran (the standard diet treatment); (ii) pieces of a high-density PE shopping bag (the PE diet treatment); (iii) pieces of styrofoam packaging (the PS diet treatment); or nothing (the starvation diet treatment). Insect colonies were maintained at room temperature and ambient moisture for 10 to 11 days, with dead insects removed every one to three days. For each cohort, gut samples for metagenomic sequencing were collected on days 10 and 11 (30 mealworms or five superworms per day). This procedure was repeated for all cohorts, yielding six replicates per insect species per treatment (three independent cohorts each with two within-cohort replicates [days 10 and 11]). See **Figure 1** for a schematic summary of the experimental design.

**Figure 1.**
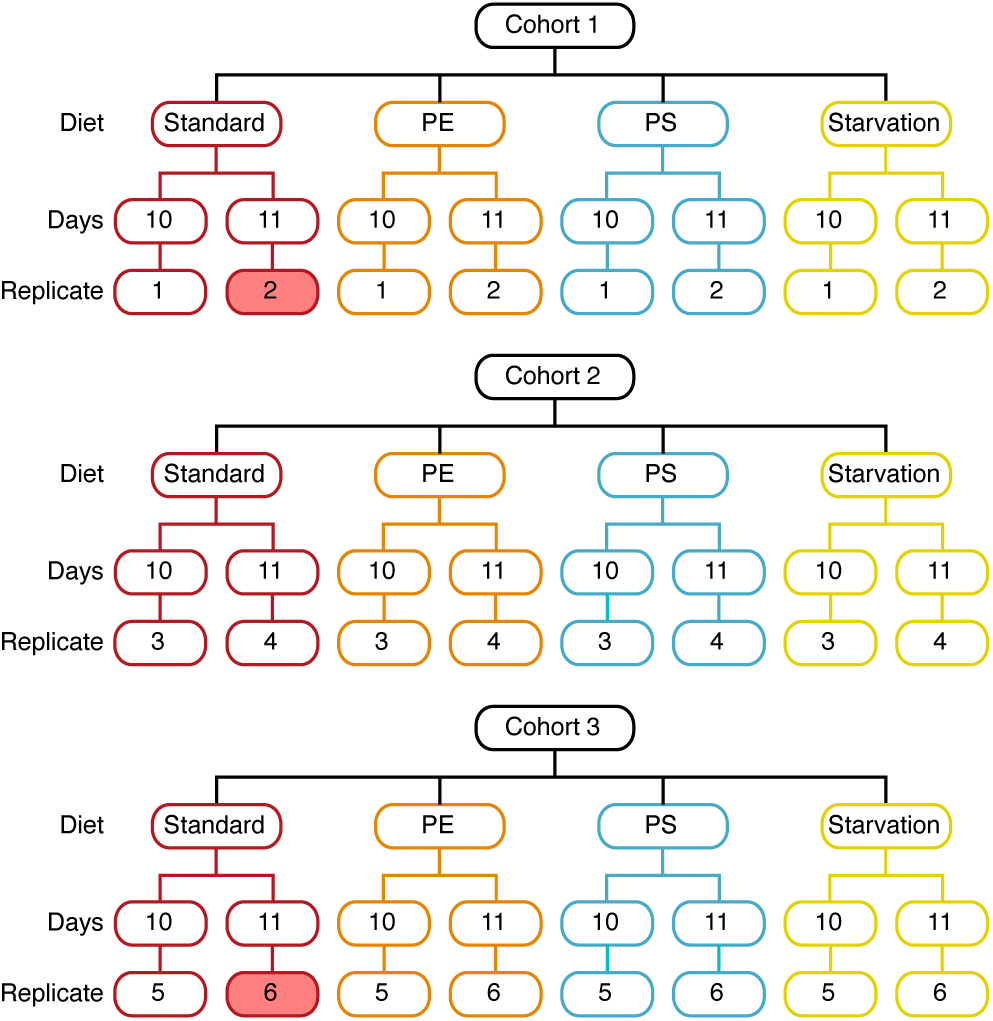
Schematic overview of the experimental design for the shotgun metagenomics. The same experimental design was replicated for mealworms and superworms. Eash insect species was purchased in three separate batches (representing distinct starting cohorts), each on a different week. For each cohort, following a two-day standardization period, insects were divided across the four diet treatments: standard (oatmeal and bran), PE (polyethylene; plastic bag), PS (polystyrene; styrofoam packaging), and starvation (no food or plastic). Following 10 or 11 days of rearing, the gut microbiota was collected from 30 (mealworms) or 5 (superworms) insects per treatment. Two mealworm samples (shown with red fill) were excluded from downstream processing as the DNA yield was too low for shotgun metagenomic library prep.

### Insect handling and DNA recovery

Each replicate consisted of the pooled gut microbial community of 30 mealworms or five superworms sampled at 10 (replicates 1, 3, 5) or 11 (replicates 2, 4, 6) days. Prior to isolation of the gut microbial communities, insects were placed in 70% ethanol, following which their exoskeleton was cut open and the digestive tract removed and placed in a sterile 0.85% NaCl saline solution. Digestive tracts were sliced and their contents squeezed into the saline. The cell suspension was transferred to a 1.7 mL tube and centrifuged at 1,000 × *g* for three minutes to remove insect remnants. The supernatant was then transferred to a fresh 1.7 mL tube and centrifuged at 16,000 × *g* for one minute to pellet the microbial cells. The pellet was resuspended in 250 µL saline and stored at 4 °C until all the samples for the day had been harvested. Total genomic DNA was isolated using ZymoBIOMICS DNA miniprep kits (Zymo Research) following the manufacturer’s protocol, with bead beating performed for 40 minutes using a Vortex Genie 2 at max speed.

### Metagenomic sequencing, assembly and binning

Illumina sequencing and library preparation were performed at Génome Québec (Montréal, QC, Canada). Briefly, libraries were generated using the NEBNext Ultra II DNA Library Prep Kit for Illumina (New England BioLabs) as per the manufacturer’s recommendations. Libraries were normalized, loaded on an Illumina NovaSeq 6000 S4 lane, and the sequencing performed for 2×150 cycles (paired-end mode). Low DNA yields precluded library preparation for two of the six mealworm standard diet replicates, and thus, these samples were excluded from downstream analysis.

Full details for the metagenomic assembly and binning are provided in **Supplementary Text S1** and are summarized below. Illumina reads were filtered using BBduk v. 38.96 [30] and trimmed using Trimmomatic v. 0.39 [31]. Trimmed reads were aligned against published mealworm or superworm genomes [32], as appropriate, using bowtie2 v. 2.4.5 [33], after which reads mapping to the insect host genomes were then discarded. Metagenomes were individually assembled for each of the 46 sequencing libraries using SPAdes V. 3.15.4 [34, 35], and scaffolds less than 2.5 kb were discarded from the assemblies using pullseq v. 1.0.2 (github.com/bcthomas/pullseq).

To facilitate the binning of scaffolds into MAGs, bowtie2 v. 2.4.1 was used to map all 22 mealworm libraries against the 22 mealworm metagenomes, and all 24 superworm libraries against all 24 superworm metagenomes. An optimized set of MAGs was produced for each of the 46 metagenomes using MetaBAT2 v. 2.14 [36], MaxBin2 v. 2.2.7 [37], CONCOCT v. 1.1.0 [38], and DAS_Tool v. 1.1.4 [39]. Finally, non-redundant sets of bacterial MAGs were separately prepared from mealworms and superworms using dRep v. 3.2.2 [40], which was parameterized to only report MAGs with at least 70% genome completeness and less than 10% genome contamination.

To identify high-quality eukaryotic bins, dRep was rerun without the completeness or contamination filters. Genome completeness metrics were then calculated for each bin using BUSCO v. 5.3.2 with the eukaryota_odb10 database [41]. Bins with at least 70% of the 255 marker genes identified as complete and single-copy, and with less than 10% of the marker genes identified as duplicated, were considered high-quality eukaryotic bins. Taxonomic classification of eukaryotic bins was then performed using BUSCO with the auto-lineage-euk function.

### Taxonomic classification and phylogenetic analysis of bacterial MAGs

MAGs were taxonomically classified using the GTDB-Tk v. 2.6.1 with the R226 database [42, 43] and the following dependencies: Prodigal v. 2.6.3 [44], pplacer v. 1.1.alpha19 [45], HMMER [46], skani v. 0.3.1 [47], and FastTree v. 2.1.11 [48]. MAGs were additionally clustered into species based on pairwise average nucleotide identity (ANI) values as calculated with FastANI v. 1.33 [49], using an ANI threshold of ∼95%.

A maximum likelihood phylogeny of all mealworm and superworm bacterial MAGs was constructed largely as described previously [50]. Briefly, homologs of the bac120 gene set were identified in all MAGs and then aligned and concatenated with the GTDB-Tk align function, after which a phylogeny was constructed using IQ-TREE2 v 2.2.2.4 [51] with the LG+F+R8 model as it was identified as the best scoring model using ModelFinder [52]. Phylogenies were visualized using iTOL [53].

### Calculations of the relative abundances of MAGs

All mealworm MAGs were combined as one multifasta file, while all superworm MAGs were combined as a second multifasta file. Next, bowtie2 v. 2.4.5 was used to map all 22 mealworm or 24 superworm Illumina sequencing libraries against the mealworm and superworm MAG multifasta files, respectively, and the output sorted with samtools v. 1.11 [54]. Custom perl scripts were then used to calculate the relative abundances of each MAG across samples either as (i) reads mapped per million reads or (ii) average sequencing depth per million reads.

### Functional annotation of metagenomes and MAGs

MAGs and metagenome assemblies (limited to scaffolds ≥2.5 kb) were functionally annotated using Distilled and Refined Annotation of Metabolism (DRAM) v. 1.2 [55] on the KBase webserver [56]. MAGs were additionally screened for potential biosynthetic gene clusters using antiSMASH v. 7.1.0 [57] with the dependencies diamond v. 2.1.8 [58], FastTree v. 2.1.11, HMMER v. 2.3.2, HMMER v. 3.4, BLAST+ v. 2.16.0 [59], and Prodigal.

### Identification and analysis of secreted proteins of interest

The 46 predicted metaproteomes (generated with DRAM) were extracted and combined into a single multifasta file. Proteins were clustered using CD-HIT v. 4.8.1 [60] to produce a non-redundant protein set, using a sequence identity threshold of ≥99% and an alignment coverage threshold of ≥99% the length of the longer sequence. The program signalP v. 6.0 [61] was then used to search for secretion tags to generate a predicted secretome. This secretome was combined with the published sequences of 12 proteins (not necessarily from insect-associated bacteria) previously implicated in plastic biodegradation [29, 62–69], as well as the published sequence of a glutathione peroxidase that lacked activity on PE in a recent study [29]. The resulting set of proteins was then searched for proteins of interest following a modified reciprocal hit approach as described in **Supplemental Text S2** using HMMER v. 3.3 and hidden Markov models (HMMs) of the Pfam database [70]. Sequence similarity networks (SSNs) were constructed using the identified proteins and the online enzyme function initiative’s enzyme similarity tool (EFI-EST; https://efi.igb.illinois.edu/efi-est/) [71, 72]; alignment score thresholds of 75, 75, and 36 were used for laccases, DyP-type peroxidases, and glutathione peroxidases, respectively. The resulting networks were visualized using Cytoscape version 3.10.1 [73]. Lastly, tBLASTn v. 2.10.1 was used to search the proteomes of the 153 dereplicated MAGs with secreted proteins of interest as queries.

### Statistical analyses

All statistical analyses were run in RStudio v. 2025.9.2.418 [74] with R v. 4.5.2 [75] and the packages ANCOMBC v. 2.12.0 [76], car v. 3.1-3 [77], dplyr v. 1.1.4 [78], emmeans v. 2.0.1 [79], ggplot2 v. 4.0.1 [80], kableExtra v. 1.4.0 [81], knitr v. 1.51 [82], lme4 v. 1.1-38 [83], patchwork v. 1.3.2 [84], phyloseq v 1.54 [85], tidyr v. 1.3.2 [86], and vegan v. 2.7-2 [87]. Full details for the statistical analysis are provided in **Supplementary Text S3** and are summarized below.

Shannon and Simpson diversity indices were estimated using the phyloseq::estimate_richness function. Statistical analysis of the log-transformed Shannon and Simpson diversity indices was performed using linear mixed models with the lmer function (lme4 package). In these models, fixed effects included insect host species, sequencing read count (millions of reads following QC), and their interaction, while insect cohort nested within species (six cohorts total, three for each species) was included as a random effect. For comparisons of diversity across diets within an insect species, diet and sequencing read count (millions of reads following QC) were included as fixed effects while insect cohort (three groups) was included as a random effect.

Bray-Curtis distances were calculated using the phyloseq::distance function. Principle Component Analyses (PCoA) and capscale analyses were run using the phyloseq::ordinate function. For analyses across insect species, PERMANOVA was run with the distance matrix as the response variable and insect species, sequencing read count (millions of reads following QC), and their interaction as main effects. For analyses across diets within an insect species, PERMANOVA and capscale analyses were run with the distance matrix as the response variable and with diet, insect cohort (three groups), and sequencing read count as main effects.

Differential abundance of taxa across diets per insect species was examined using ANCOM-BC2. ANCOM-BC2 was run with a model that included diet as a fixed effect and insect cohort (three levels) as a random effect. P value adjustment was performed using Benjamini-Hochberg corrections. In addition, ANCOM-BC2 was run twice per dataset, once with the standard diet as the reference condition and once with the starvation diet as the reference condition. A taxon was considered differentially abundant if, in both comparisons, the adjusted p-value was ≤ 0.025 and the taxon passed the sensitivity analysis.

## Data availability

Raw sequencing data, assembled metagenomes, and MAGs were deposited in the National Center for Biotechnology Information (NCBI) database under BioProject accession PRJNA1399388. Code to repeat all computational analyses described in this manuscript is available via GitHub at github.com/diCenzo-Lab/018_2026_Beetle_PE_PS_metagenomics.

## RESULTS

### Overview of the experimental design

We used a metagenomic approach to: (i) study the biodiversity and functional potential of the gut microbiota of mealworms and superworms, (ii) characterize the impacts of plastic ingestion on the gut microbiome, and (iii) identify enzymes of potential relevance for plastic biodegradation. Due to the low biomass obtained from the guts of individual insects, each replicate consisted of the pooled microbiota of either 30 mealworms or 5 superworms. In addition, the time required to dissect the insects necessitated that each set of replicates be collected on separate days. As a result, insects were purchased in three batches on separate weeks (representing distinct starting cohorts), and two replicates collected per cohort on days 10 and 11 of rearing (**Figure 1**). Consequently, for each insect species, six replicates per treatment were collected, with pairs of replicates nested within three insect cohorts. The exception was for mealworms reared with a standard diet, as two replicates (one from each of insect cohorts 1 and 3) were lost due to insufficient DNA yields.

### Summary statistics of the metagenomic sequencing

Between 52 million and 79 million read pairs (16 to 24 Gb, median of 19 Gb) were generated per sample (**Dataset S1**). Following trimming and removal of reads mapping to the insect host genome, between 15 million and 68 million read pairs (4.6 to 20 Gb, median of 15 Gb) and between 18 million and 59 million read pairs (5.6 to 18 Gb, median of 12 Gb) remained for the mealworm and superworm samples, respectively (**Dataset S1**). Following assembly, the resulting metagenomes ranged in size from 53 Mb to 436 Mb (**Dataset S1**); after removal of scaffolds less than 2.5 kb in length, the metagenomes were reduced by 17% to 65% and retained between 34 Mb to 204 Mb (**Dataset S1**). The superworm metagenomes (307 Mb ± 68 Mb; or 149 Mb ± 35 Mb following removal of scaffolds < 2.5 kb) were on average larger than the mealworm metagenomes (117 Mb ± 29 Mb; or 74 Mb ± 19 Mb following removal of scaffolds < 2.5 kb).

Binning resulted in the production of 9 to 36 MAGs per assembly, with a range of 9 to 23 (median 18) MAGs per mealworm assembly and 10 to 36 (median 25) MAGs per superworm assembly. MAGs were separately dereplicated for the mealworm and superworm samples with an ANI threshold of 99%, resulting in 53 and 100 high-quality bacterial MAGs (≥70% completeness and ≤10% contamination; median values of 98.9% completeness and 0.9% contamination) for the mealworm and superworm samples, respectively (**Dataset S2**). The 53 mealworm MAGs represent 44 species across 28 genera and four phyla, while the 100 superworm MAGs represent 88 species across 47 genera and five phyla (**Figure 2, Dataset S2**). Unexpectedly, only 19 of the species (from 13 genera) were common between the two insect hosts. Overall, the 153 MAGs represent 113 species, including 45 putatively novel species and one previously undescribed family (**Figure 2, Dataset S2**). In addition, one high-quality eukaryotic bin was obtained from the mealworm samples and assigned to the class Saccharomycetes by BUSCO (96.9% complete single copy, 0.2% complete duplicated BUSCO saccharomycetes_odb10 genes). No archaeal MAGs were recovered.

**Figure 2.**
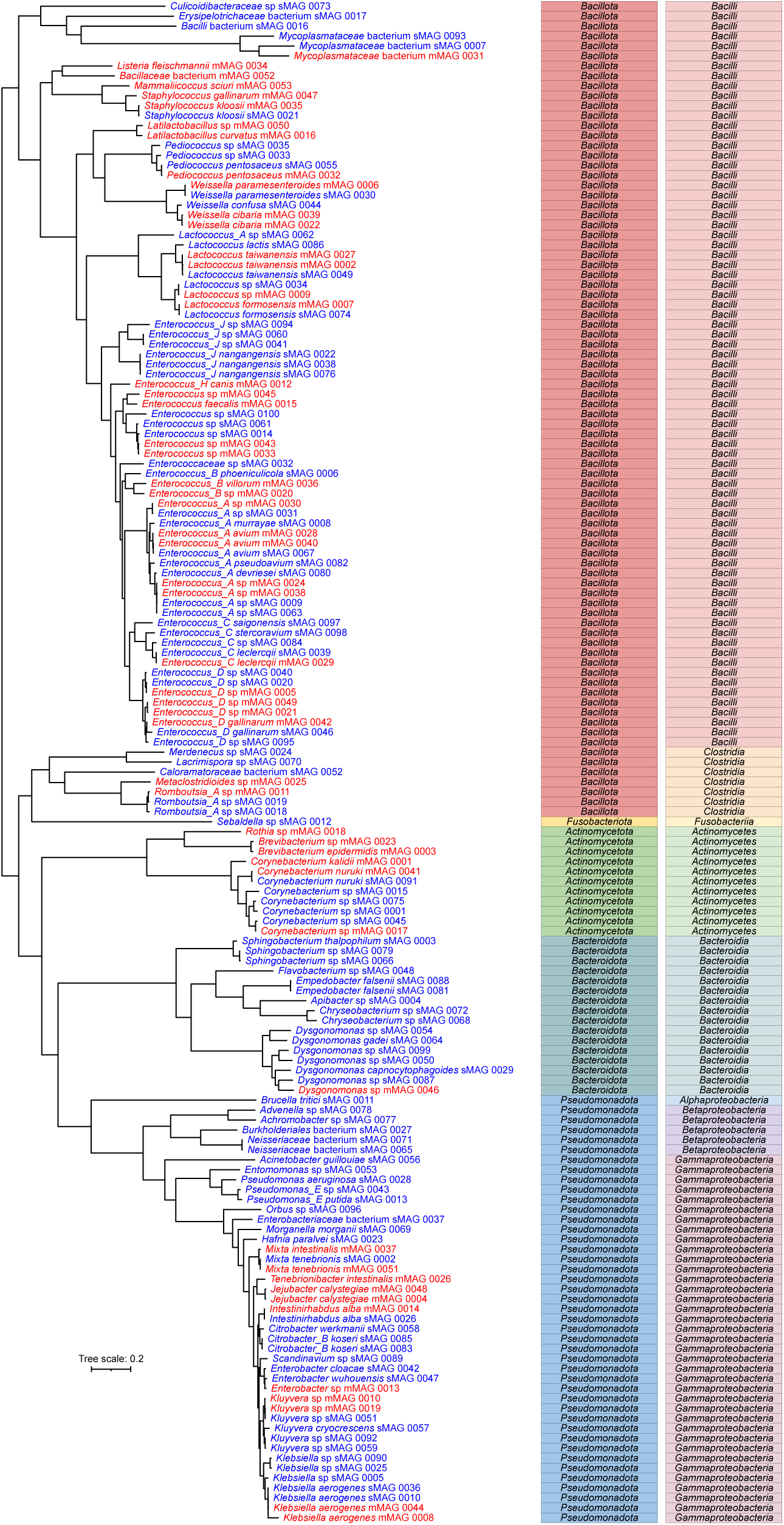
Diversity of the metagenome-assembled genomes (MAGs) recovered from mealworm and superworm gut metagenomic datasets. An unrooted, maximum-likelihood phylogeny of the 53 dereplicated and high-quality mealworm MAGs (red) and the 100 dereplicated and high-quality superworm MAGs (blue). The columns to the right of the phylogeny indicate the taxonomic classification of each MAG at the level of phylum (left) or class (right). The scale bar represents the average number of amino acid substitutions per site. An interactive version of this phylogeny with support values (SH-aLRT values provided as bootstrap0 and ultra-fast jackknife values as boostrap1) is available at itol.embl.de/shared/qiBXfvsqoMD0.

### Mealworm and superworm gut microbiomes were dominated by bacteria

As noted above, following dereplication, 53 and 100 bacterial MAGs were recovered from the mealworm and superworm metagenomics datasets, respectively, whereas only a single, high-quality fungal bin (a yeast) was recovered. An average of 90% (standard deviation of 3.6%) of reads from the mealworm datasets mapped to the mealworm bacterial MAGs, while an average of 71% (standard deviation of 9.0%) of reads from the superworm datasets mapped to the superworm bacterial MAGs (**Dataset S1**). The lower rate of mapping for the superworm datasets may reflect a higher abundance of insect host reads in the final Illumina datasets (**Supplementary Text S4**) rather than due to a lower rate of MAG recovery. Regardless, these results indicate the MAGs captured most of the available gut microbiome information, and they are consistent with most of the sequencing reads corresponding to bacterial DNA. The mapping also indicates that the lack of fungal bins reflects a low abundance of fungal DNA in the datasets rather than poor recovery of fungal bins from the assemblies, as also supported by colony forming unit counts (**Supplementary Text S5 and Table S1**).

### Low complexity and high biosynthetic potential of mealworm and superworm gut microbiomes

To analyze the composition of the mealworm and superworm gut bacterial communities, the shotgun metagenomics data were mapped to the corresponding set of dereplicated bacterial MAGs, and the abundance of each MAG was calculated for each sample. Initially, we wished to better understand the generic composition of the bacterial communities of mealworms and superworms, and how they differ, and thus we began by comparing the communities between insects.

The Shannon and Simpson diversity indices trended towards being 1.39-fold (95% confidence interval of 0.91-fold to 2.13-fold) and 1.17-fold (95% confidence interval of 0.83-fold to 1.64-fold) higher, respectively, for the superworm gut bacterial communities compared to the mealworm gut bacterial communities, although neither comparison was statistically significant (**Figure 3**). Higher diversity in the superworm gut communities is further supported by the observation that the assembled superworm metagenomes were on average larger than the mealworm metagenomes, as noted above, with a higher number of MAGs recovered from the superworm data compared to the mealworm data. Moreover, the 10 most abundant MAGs in any given sample accounted for an average of 75% the total relative abundance of the MAGs in the superworm communities but 89% in the mealworm communities.

**Figure 3.**
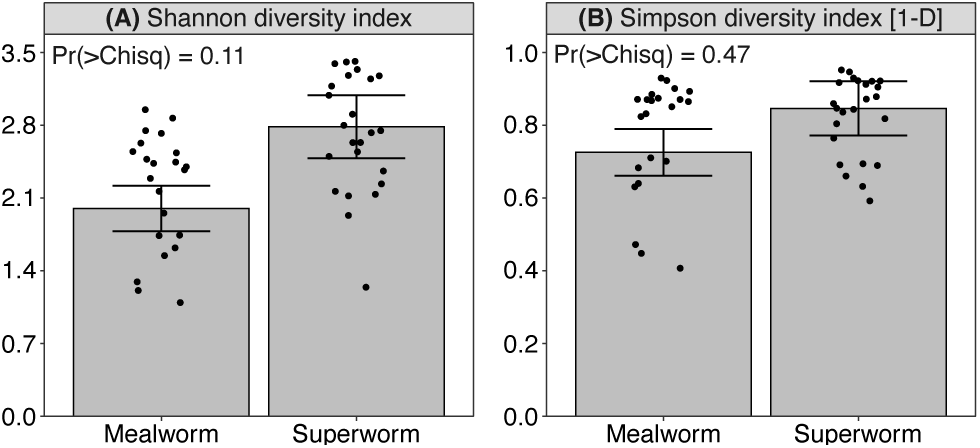
Alpha-diversity indices for mealworm and superworm gut bacterial communities. (**A**) Shannon and (**B**) Simpson diversity indices are shown. The dots show the raw values for individual samples, while the bars and the error bars show the estimated marginal means and their standard errors. The p-values are from Chi-squared tests generated from running Type II ANOVAs on mixed linear models that included insect species, read depth, and their interaction, as fixed effects, and insect cohort as a random effect.

The mealworm and superworm gut bacterial communities were significantly different at all taxonomic levels (**Figure S1**), with the strongest effect at the taxonomic ranks of species (PERMANOVA, Pr(>F) = 0.0001; 42% of the total variation was explained by insect species) and genus (PERMANOVA, Pr(>F) = 0.0001; 31% of the total variation was explained by insect species). Notably, the genera *Citrobacter* and *Entomomonas*, as well as the unnamed genera *Citrobacter_B* (family *Enterobacteriaceae*), *CALYQQ01* (family *Enterobacteriaceae*), and *JAGNPU01* (family *Neisseriaceae*) had relative abundances of ≥ 1% across the superworm gut communities but were absent from all mealworm gut communities, while the genera *Jejubacter*, *Latilactobacillus*, and *Tenebrionibacter* had relative abundances of ≥ 1% across the mealworm gut communities but were absent from all superworm gut communities (Table 1).

**Table 1.** Composition of the mealworm and superworm gut bacterial communities.

| Taxon | Mealworm | Superworm |
| --- | --- | --- |
| **Phyla** |  |  |
| *Actinomycetota* | 1.9 ± 3.4 | 1.0 ± 0.8 |
| *Bacillota* | 63 ± 24 | 65 ± 20 |
| *Bacteroidota* | 0.1 ± 0.5 | 5.3 ± 2.9 |
| *Fusobacteriota* | 0.0 ± 0.0 | 0.3 ± 0.3 |
| *Pseudomonadota* | 35 ± 23 | 28 ± 18 |
| **Genera** |  |  |
| CALYQQ01 (family *Enterobacteriaceae*) | 0.0 ± 0.0 | 1.9 ± 1.1 |
| *Citrobacter* | 0.0 ± 0.0 | 2.8 ± 2.3 |
| *Citrobacter_B* (family *Enterobacteriaceae*) | 0.0 ± 0.0 | 2.2 ± 3.5 |
| *Entomomonas* | 0.0 ± 0.0 | 3.6 ± 5.0 |
| JAGNPU01 (family *Neisseriaceae*) | 0.0 ± 0.0 | 1.0 ± 0.6 |
| *Jejubacter* | 8.9 ± 9.6 | 0.0 ± 0.0 |
| *Latilactobacillus* | 6.6 ± 7.3 | 0.0 ± 0.0 |
| *Tenebrionibacter* | 8.1 ± 11 | 0.0 ± 0.0 |

\* All phyla detected in at least one insect species are provided, while the listed genera are limited to those with an average relative abundance of ≥ 1% in one insect species but absent in the other insect species.

On the other hand, the mealworm and superworm gut communities were only weakly separated from each other based on Bray-Curtis distances at the phylum level (PERMANOVA, Pr(>F) = 0.056; 5.7% of the total variation was explained by insect species) (**Figure S1F**). Indeed, the gut bacterial communities of mealworms and superworms were both dominated by the phyla *Bacillota* and *Pseudomonadota*, which collectively accounted for an average of > 90% of these communities (Table 1). The phyla *Actinomycetota* and *Bacteroidota* were also found as minor components across both insect species, while the superworm gut microbiota additionally included a low abundance of the phylum *Fusobacteriota* (Table 1).

The biosynthetic potential of the bacterial component of the mealworm and superworm gut microbiota was investigated using antiSMASH. Overall, 132 of the 153 mealworm or superworm MAGs carried at least one biosynthetic gene cluster (BGC) (**Dataset S3**). These 132 MAGs collectively carried at least 451 BGCs, with a median of 3 BGCs per MAG (range of 1 to 11). The identified BGCs include 70 encoding non-ribosomal peptide synthases (NRPSs), 100 poly keytide synthases (PKSs), 167 ribosomally synthesized and post-translationally modified peptides (RiPPs), 40 terpenes, and 74 BGCs encoding other products (**Dataset S3**).

### Plastic ingestion had little impact on the mealworm and superworm gut bacterial communities

We next explored whether ingesting plastics impacted the gut microbiomes of mealworms or superworms. We therefore compared the gut bacterial communities of insects fed a standard diet of oatmeal and bran to those fed only PE or only PS, as well as those provided no food to control for the effects of starvation. Plastic ingestion had no statistically significant impact on the overall bacterial community alpha diversity metrics in either insect (**Figures 4A-B, 4D-E**). However, we note that with one exception (mealworms provided PS), the estimated marginal means for the Shannon diversity indices for insects provided plastics or a starvation diet were higher than those provided oatmeal and bran (**Figures 4A, 4D**), suggesting that starvation may have resulted in a slight (but not statistically significant) increase in overall diversity of the mealworm and superworm gut microbiota.

**Figure 4.**
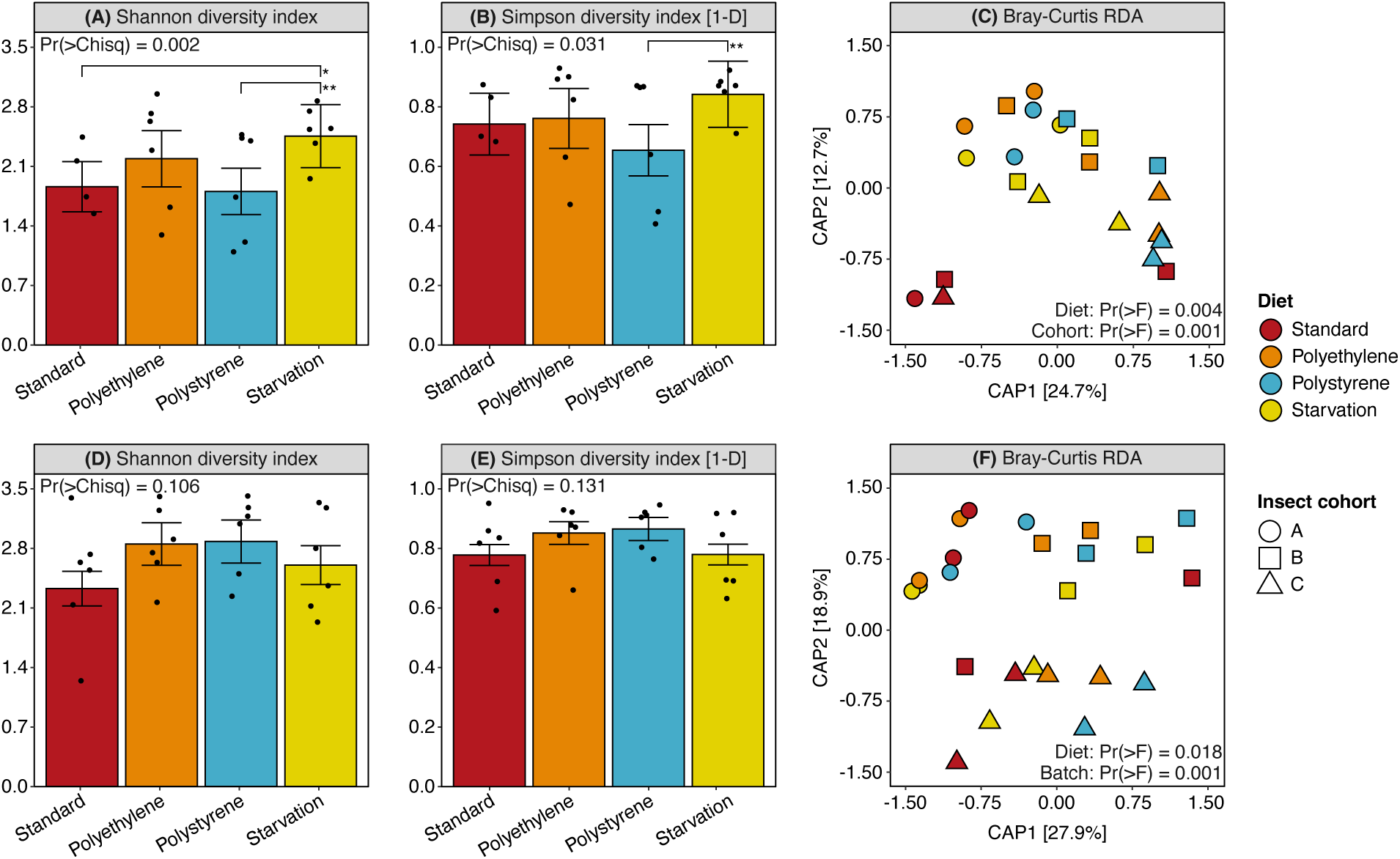
Diversity metrics of mealworm and superworm gut bacterial communities fed different diets. Results from alpha-diversity and beta-diversity analyses are shown for (**A**-**C**) mealworm and (**D**-**F**) superworm gut bacterial communities. For each insect species, the panels show (**A**, **D**) Shannon and (**B**, **E**) Simpson diversity indices, as well as (**C**, **F**) an ordination plot produced from a capscale analysis run on Bray-Curtis distances. (**A**, **B**, **D**, **E**) For the Shannon and Simpson diversity indices, the dots show the calculated metrics for individual samples, while the bars and the error bars show the estimated marginal means and their standard errors. The p-values are from Chi-squared tests generated from running Type II ANOVAs on mixed linear models that included diet and sequencing depth as fixed effects and insect cohort as a random effect. The brackets represent emmeans contrasts with a p-value < 0.1 (*) or 0.05 (**). (**C**, **F**) For the ordination plots, p-values are from F-tests generated from running Type II ANOVAs on capscale analyses run with models that included diet, sequencing depth, and insect cohort as main effects.

A capscale redundancy analysis (RDA) found statistically significant effects of diet on the gut bacterial communities in both mealworms (Pr(>F) = 0.01; 23% of the total variation was explained by diet) and superworms (Pr(>F) = 0.025; 14% of the total variation was explained by diet). Plotting the capscale results as an ordination plot revealed that the gut bacterial communities of mealworms fed a standard diet of oatmeal and bran tended to group separately from those provided PE, PS, or nothing (**Figure 4C**). No clear separation was observed between PE, PS, and the starvation control for mealworms (**Figure 4C**). In the case of superworms, no obvious effects of diet were observable in an ordination plot (**Figure 4F**). On the other hand, clear effects of insect cohort were observable in the superworm ordination plot (**Figure 4F**), and to a lesser extent, the mealworm ordination plot (**Figure 4C**). Indeed, an ANOVA on the capscale results indicated that insect cohort had strong and statistically significant effects on the composition of the gut bacterial communities of both mealworms (Pr(>F) = 0.008; 20% of the total variation was explained by insect cohort) and superworms (Pr(>F) = 0.001; 33% of the total variation was explained by insect cohort). The same overall patterns were observed if the community abundance data were first summarized at the genus level (**Figure S2**); diet explained 19% (Pr(>F) = 0.034) and 13% (Pr(>F) = 0.075) of the total genus-level variation in the mealworm and superworm gut bacterial communities, respectively, while insect cohort explained 21% (Pr(>F) = 0.003) and 30% (Pr(>F) = 0.001).

We hypothesized that bacteria capable of degrading and catabolizing PE or PS would increase in relative abundance in the PE or PS fed insects. We therefore compared the abundance of MAGs or species in the PE or PS fed insects to insects fed a standard diet and the starved insects. In both mealworms and superworms, no MAGs or species showed a statistically significant increase in relative abundance in the PE or PS diets compared to both the standard and starvation diets.

### Enzymes of potential relevance for PE and PS degradation

We reasoned that if the gut microbiota of mealworms and superworms contained microbes capable of degrading PE or PS, the associated genes should be detectable in the metagenomes even if the associated organism(s) did not increase in abundance upon PE or PS ingestion. We therefore annotated all 46 assembled but unbinned metagenomes, resulting in a total of ∼ 5.2 million proteins across all metagenomes. We then dereplicated (≥ 99% identity over ≥ 99% the length of the longer protein) the proteins, yielding a dereplicated metaproteome of 1,318,948 proteins. Lastly, we reasoned that since plastic polymers are too large to be imported into the cell, any bacterially encoded enzymes able to modify plastics must be secreted. We therefore used signalP to predict which proteins were secreted, resulting in a final secretome of 105,322 proteins. Using HMMs, we identified numerous proteins belonging to enzyme classes previously implicated in early steps of PE or PS degradation [29, 62–69] including 186 laccases, 22 glutathione peroxidases, 99 DyP-type peroxidases, one 2OG-Fe(II) oxygenase, and one fatty acid desaturase (**Dataset S4**).

We next constructed sequence similarity networks (SSNs) for the laccases, glutathione peroxidases, and the DyP-type peroxidases, together with reference proteins previously implicated in PE or PS degradation. At the selected clustering threshold, none of the laccases clustered with the four laccases previously implicated in oxidation of PE or PS (**Figure 5A**), and thus this analysis did not provide clear support for any of the identified laccases having activity on PE. Seventeen of the 22 putative glutathione peroxidases clustered with a glutathione peroxidase previously suggested to oxidize PE/PS, with all nodes having a percent identity of ≥ 40% (**Figure 5B**). However, this cluster also included a glutathione peroxidase for which a previous study found no evidence of activity on PE [29], raising doubts as to whether any of the enzymes of this cluster would be active on PE. Lastly, only a one protein coding gene was identified for each of the 2OG-Fe(II) oxygenase family and the fatty acid desaturase family, with these genes each found in only one metagenome. The low diversity and prevalence of these enzyme classes suggest that they are unlikely to be involved in degrading PE or PS in the guts of mealworms or superworms.

**Figure 5.**
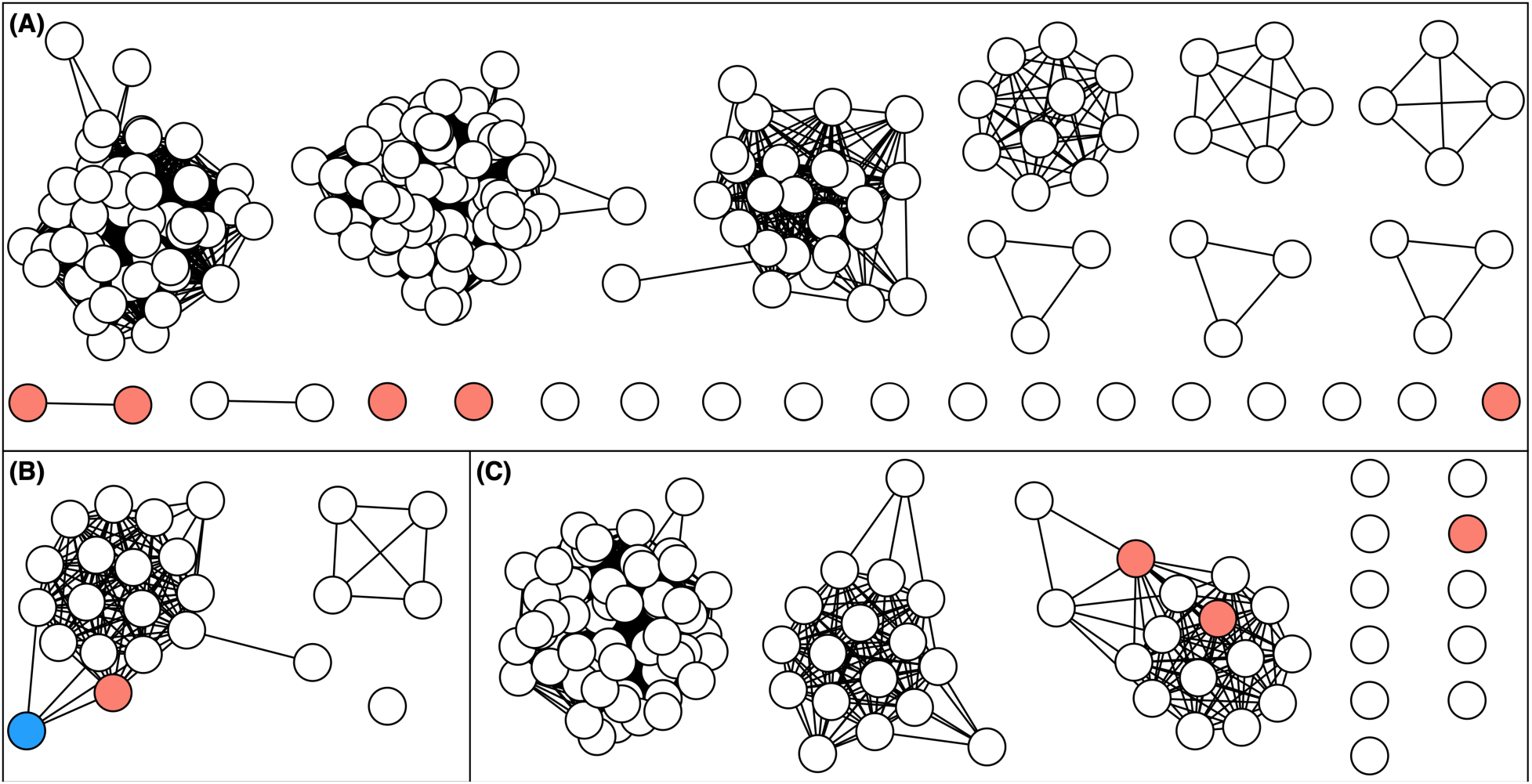
Sequence similarity networks of secreted oxidative proteins. Sequence similarity networks, calculated using EFI-EST, are shown for (**A**) laccases, (**B**) glutathione peroxidases, and (**C**) DyP-type peroxidases. Each node (the circles) represents one protein, while edges (the lines) represent sequence similarity between pairs of proteins above the threshold. White nodes represent proteins identified in the secretome of the mealworm and superworm gut microbiomes. Red notes represent proteins implicated in the biodegradation of polyethylene or polystyrene in previous studies, while the blue node represents a protein that lacked activity on polyethylene.

On the other hand, 14 of the 99 putative DyP-type peroxidases clustered with two DyP-type peroxidases previously suggested to oxidize PE in independent studies, with all nodes having a percent identity of ≥ 40% (**Figure 5C**). Mapping these 14 DyP-type peroxidases to the MAGs revealed hits with percent identities ≥ 99% for 13 of the proteins. These 13 proteins mapped to eight distinct MAGs all of which belong to the phylum *Actinomycetota*; the majority (6 of 8) of the MAGs were classified as belonging to the genus *Corynebacterium*, with the other two MAGs belonging to the genera *Brevibacterium* and *Rothia*. The remaining 58 DyP-type peroxidases not included in this primary cluster but that could be confidently assigned to a MAG, mapped to 28 unique MAGs from 13 genera across the families *Enterobacteriaceae* (18 MAGs), *Mycobacteriaceae* (6 MAGs), *Burkholderiaceae* (1 MAG), *Neisseriaceae* (1 MAG), *Bartonellaceae* (1 MAG), and *Staphylococcaceae* (1 MAG) (**Dataset S4**). Interestingly, many of the MAGs encoding DyP-type peroxidases were from lineages common to both mealworms and superworms. Indeed, MAGs from six of the 19 bacterial species shared between the insect species encode DyP-type peroxidases. In contrast, none of the MAGs encoding glutathione peroxidases belonged to a species common to both mealworms and superworms.

## DISCUSSION

Mealworms and superworms are of scientific interest in part due to their potential for hosting gut microbiota that biodegrade diverse plastics [12], although other studies report conflicting results [20]. However, most studies exploring the gut microbiome of these insects have relied on 16S rRNA amplicon sequencing. We are aware of only one study reporting high-quality, assembled metagenomes for mealworms [29] and none for superworms, and we are unaware of high-quality studies using shotgun metagenomics to explore the impact of plastic ingestion on the composition of the gut microbiomes of these insects. To address these knowledge gaps, we generated shotgun metagenomics data for mealworms and superworms with or without exposure to plastics to generate high-quality gut metagenomes for both insects. From this baseline, we further assembled a set of reference MAGs, and used them to explore the impact of ingesting PE or PS on the composition of the mealworm and superworm gut microbiomes.

### Composition of the gut microbiome of mealworms and superworms

Our metagenomics data and direct microbe isolation experiments (**Supplementary Text S5**) suggested that the gut microbiomes of mealworms and superworms are dominated by bacteria with only a low abundance of yeasts and other fungi. We found this surprising given past studies showing various beetle species collectively host diverse communities of yeasts in their guts [88–92]. These observations are not necessarily in conflict, however, as it is possible that beetle gut microbiomes contain diverse yeasts but that their absolute abundance is low in comparison to the absolute abundance of bacteria. We also cannot rule out that the abundance of fungi in the gut microbiome of industrially-reared beetle larvae would differ from beetles collected from their natural habitat. That said, the sole fungal bin that we recovered belonged to the class Saccharomycetes (division: Ascomycota), consistent with Ascomycota often being the dominant fungal component of beetle microbiomes [88–91].

In comparing the gut bacterial communities of mealworms and superworms, we found a trend, albeit not statistically significant, towards the superworm gut communities being more diverse than the mealworm gut communities. This result is consistent with studies suggesting a positive correlation between host body size and gut microbiome diversity [93–96]. At higher taxonomic levels, the mealworm and superworm gut bacterial communities were broadly similar, with both being dominated by the phyla *Bacillota* (syn. *Firmicutes*) and *Pseudomonadota* (syn. *Proteobacteria*). On the other hand, we were surprised by the proportion of taxa that were specific to mealworms or superworms at lower taxonomic levels, although we expect the overlap to increase as the gut microbiome of additional cohorts are sequenced. The dominance of the phyla *Bacillota* and *Pseudomonadota* is consistent with other studies of mealworms [11, 97–99] and superworms [19, 26, 100]. Likewise, *Pseudomonadota* tends to be a dominant member of the gut microbiome of beetle species more broadly, while a dominance of the phylum *Bacillota* in other beetle species is less common [89–91, 101, 102].

### Impact of plastics on the gut bacterial communities of mealworms and superworms

One of our objectives was to explore the impacts of PE and PS ingestion on the gut microbial communities of mealworms and superworms, reasoning that if these communities harbour microbes able to breakdown and catabolize these polymers, these microbes should increase in abundance when exposed to PE or PS. Although there was a statistically significant effect of diet on the overall composition of the gut bacterial communities of mealworms and superworms, we were unable to observe clustering of the PE or PS diets away from both the standard diet and starvation diet controls in capscale analyses. In addition, no MAGs were identified as statistically enriched in the PE or PS diets relative to both the standard and starvation diets in either insect. This contrasts with numerous other studies reporting that feeding of plastic resulted in clear changes in the gut microbiome composition of these insect species (e.g., [15–17, 19, 103]). We hypothesize that this discrepancy can be largely explained by three experimental parameters as outlined below.

In our experiments, insects were exposed to plastics for 10 days, whereas past studies exposed their insects to plastics for 14 to 45 days [11, 15–17, 19, 103, 104], while at least one study spanned multiple insect generations when the plastics were co-fed with nutritious food [105]. As such, it may be that our rearing period was not sufficiently long for the diets to have had pronounced effects on the gut microbiome. However, most mealworm samples fed a standard diet grouped separate from the rest of the mealworm samples, demonstrating that 10 days was sufficient for at least some effects of diet to be observable. As such, the length of the experiment cannot fully explain the lack of easily observable effects of PE or PS ingestion on the gut bacterial communities of mealworms and superworms.

In designing our experiments, we chose to include a treatment wherein the insects remained unfed, provided no food or plastic, to control for the effects of starvation on the gut microbial communities. This control proved to be important. Several differences were detected between the gut bacterial communities of insects provided PE or PS compared to the standard diets; however, these same differences were not observed when PE- or PS-fed insects were compared with starved controls. In contrast to our study, with a few exceptions (e.g., [25, 26]), most previous studies exploring the effect of plastics on the gut microbiome of mealworms or superworms have not included a starvation treatment to control for the effects of nutrient deprivation. As such, we hypothesize that many of the taxa previously reported to respond to plastic exposure in the guts of these insects can potentially be explained by starvation rather than a direct impact of plastics.

Lastly, our experimental design involved obtaining insects from a local pet store in three batches, representing three distinct cohorts. The redundancy analysis revealed that the insect cohort had a strong impact on the gut microbial community, with cohort explaining more of the total variation in community composition than diet. These results suggest that each cohort of mealworms or superworms had very different starting gut microbiomes and that the diet treatments did not result in their convergence. In addition, there was clear grouping of the samples based on insect cohort in the ordination plots, leading us to hypothesize that the effect of cohort may have masked any potential effects of diet. This strong effect of cohort has interesting implications to consider in future studies. First, it highlights that, unsurprisingly [11], the gut microbiomes of mealworms and superworms differ between cohorts, perhaps due to variations in age, health, environment, or diet prior to purchase [95, 106, 107]. Second, these results suggest that investigations of the effect of diet (or other factors) on the gut microbiomes of these insects should at least initially be performed using replicates that all come from a common starting cohort. Doing so would provide more power to detect diet-driven effects on the gut microbiome of a given cohort by ensuring the starting microbiome of each sub-cohort is similar. However, doing so comes with the important caveat that the results may not be generalizable to other cohorts of the same species.

### DyP-type peroxidases as candidate biocatalysts for oxidation of polyethylene

Initially, we wished to identify MAGs enriched in the plastic diet treatments to limit our search for PE/PS modifying enzymes to the enzymes encoded by these MAGs. However, given the lack of enrichment of MAGs in these treatments, we instead took an untargeted approach and searched the entire predicted metaproteomes generated from annotating the 46 assembled metagenomes. While the biodegradation of PE and PS likely requires a multi-step pathway [108, 109], here we focused just on enzymes that could potentially catalyze early PE/PS modification, and specifically, oxidative enzymes from enzyme classes previously implicated in biodegradation of PE/PS including laccases [64, 67–69], glutathione peroxidases [64], and DyP-type peroxidases [29, 63]. We additionally focused on secreted enzymes given that plastic polymers are unlikely to be imported into a bacterial cell and thus only enzymes secreted outside of the cell would have access to the plastic.

While we identified many secreted enzymes with similarity to laccases, none grouped with laccases previously suggested to be involved in PE degradation, and thus we have low confidence that laccases are an important contributor to PE degradation in the mealworm and superworm gut environments. Similarly, we identified numerous putative glutathione peroxidases amongst the secreted proteins that clustered with an enzyme previously suggested to oxidize PE [64]. However, this protein cluster also included an enzyme that had no detectable activity with PE as a substrate in a recent study [29], giving us low confidence that any of the enzymes in this cluster are active on PE.

We also identified 99 secreted enzymes showing similarity to DyP-type peroxidases. Fourteen of these enzymes clustered with two DyP-type peroxidases with published evidence of activity on PE [29] or pre-oxidized PS [63]. Twelve of these 14 enzymes come from *Corynebacterium* spp. as does one of the literature enzymes [29], whereas the other literature enzyme comes from *Thermomonospora curvata* B9^T^ that was originally isolated from straw [63, 110]. Notably, experimental evidence suggests that at least some *Corynebacterium* isolates can oxidize PE [29], and in our study, *Corynebacterium* was one of the 14 genera for which we recovered MAGs from both mealworms and superworms, with *Corynebacterium* MAGs recovered from both insect species found to encode DyP-type peroxidases. More broadly, we see a slight enrichment of DyP-type peroxidases in taxa common between the mealworm and superworm MAG sets (46% of genera and 32% of species) relative to taxa identified in only one insect host (23% of genera and 20% of species). Considering that the ability to degrade PE/PS is likely to be rare in nature, we consider it more parsimonious if any PE/PS-degrading ability of the gut microbiomes of mealworms and superworms is due to the taxa common to both insect hosts rather than distinct taxa. Consequently, we interpret our results (SSN clustering and broad distribution of DyP-type peroxidases amongst the shared microbial taxa), together with recent experimental data on PE [29] and pre-treated PS [63], as suggesting that DyP-type peroxidases are likely to be the dominant enzyme responsible for PE/PS oxidation in the gut environments of mealworms and superworms, should it be occurring.

## ACKNOWLEDGEMENTS

We thank Philippe Daoust and Sharen Roland and other personnel at Génome Québec for the advice on sequencing strategies and for performing all the Illumina sequencing reported in this study. This research was supported by the “Optimizing a microbial platform to break down and valorize waste plastic” project funded by the Government of Canada through Genome Canada and Ontario Genomics (OGI-207), the Government of Ontario through an Ontario Research Fund (ORF)—Large Scale Applied Research Project (LSARP) grant (File 18414), and Imperial Oil Limited through a University Research Award. Additionally, this research was enabled, in part, through computational resources provided by Compute Ontario (computeontario.ca) and Digital Research Alliance of Canada (alliancecan.ca). LAH was supported by the Canada Research Chairs.

