## Supplementary Materials for "Host cohort has a larger impact on the gut microbiomes of mealworms and superworms than does the ingestion of polyethylene or polystyrene"

**This PDF file includes:**

Text S1-S5 (pages 2-5)
Table S1 (page 6)
Figures S1-S2 (pages 7-8)
Legends for Datasets S1-S4 (page 9)
Supplementary References (page 10)

**Other supplementary materials for this manuscript include:**

Datasets S1-S4 (xlsx)

### SUPPLEMENTAL TEXT

#### Text S1. Metagenomic sequencing, assembly and binning

Illumina sequencing and library preparation were performed at Génome Québec (Montréal, QC, Canada). Briefly, libraries were generated using the NEBNext Ultra II DNA Library Prep Kit for Illumina (New England BioLabs) as per the manufacturer's recommendations. Libraries were normalized, loaded on an Illumina NovaSeq 6000 S4 lane, and the sequencing performed for 2x150 cycles (paired-end mode). Low DNA yields precluded library preparation for two of the six mealworm standard diet replicates, and thus, these samples were excluded from downstream analysis.

Illumina reads were filtered using BBduk v. 38.96 and trimmed using Trimmomatic v. 0.39 with the parameters: LEADING:3 TRAILING:3 SLIDINGWINDOW:4:15 MINLEN:36. Trimmed reads were then aligned against previously published mealworm or superworm genomes, as appropriate, using bowtie2 v. 2.4.5. Reads mapping to the insect host genomes were then discarded. Subsequently, SPAdes V. 3.15.4 was used to individually assemble each of the 46 sequencing libraries into metagenomes using the meta and only-assembler flags, and *k*-mer sizes of 33, 55, 77, 99, and 127. Scaffolds less than 2.5 kb were discarded from the assemblies using pullseq v. 1.0.2.

To facilitate the binning of scaffolds into MAGs, an all-against-all mapping of the Illumina reads for the microbiomes associated with each insect species was performed using bowtie2 v. 2.4.1 and sorted with samtools v. 1.11; i.e., all 22 mealworm libraries were mapped against all 22 mealworm metagenomes, and all 24 superworm libraries were mapped against all 24 superworm metagenomes. The metagenome scaffolds and abundance data were then binned into MAGs using MetaBAT2 v. 2.14, MaxBin2 v. 2.2.7, and CONCOCT v. 1.1.0 using default parameters, except that a chunk size of 10,000 and an overlap size of 0 were used with CONCOCT. For each of the 46 metagenomes, an optimized set of MAGs was produced using DAS_Tool v. 1.1.4 with DIAMOND v. 2.0.14 and the output of the three binning algorithms. Finally, non-redundant sets of bacterial MAGs were separately prepared from the mealworm and the superworm DAS_Tool outputs using dRep v. 3.2.2 and the following dependencies: Mash v. 2.3 [1], MUMmer v. 4.0.0rc1 [2], CheckM v. 1.1.3 [3], Prodigal v. 2.6.3, and pplacer v. 1.1.alpha19. dRep was parameterized to only report MAGs with at least 70% genome completeness and less than 10% genome contamination, and it was run once with the 22 sets of mealworm MAGs as input and separately with the 24 sets of superworm MAGs as input. The quality of the final sets of MAGs was confirmed using CheckM v. 1.2.3 run in the lineage_wf mode.

To identify high-quality eukaryotic bins, dRep was rerun without the completeness or contamination filters. Genome completeness metrics were then calculated for each of the resulting bins using BUSCO v. 5.3.2 with the eukaryota_odb10 database, with the dependencies MetaEuk v. 1da320a [4], HMMER v. 3.3, and BLAST+ v. 2.10.1. Bins with at least 70% of the 255 marker genes identified as complete and single-copy, and with less than 10% of the marker genes identified as duplicated, were considered high-quality eukaryotic bins. Taxonomic classification of eukaryotic bins was then performed using BUSCO with the auto-lineage-euk function.

#### Text S2. Modified reciprocal hit approach for protein identification

Our protein database was searched following a modified reciprocal hit approach. First, the proteins were searched using the HMMSEARCH function of HMMER v. 3.3 and hidden Markov models (HMMs) corresponding to five protein families putatively associated with plastic degradation: laccases (Pfam families PF00394, PF07731, and PF07732), glutathione peroxidases (PF00255), DyP-type peroxidases (PF04261, PF21105, and PF20628), 2OG-Fe(II) oxygenases (PF13532), and fatty acid desaturases (PF00487). The HMMs were created using the HMMBUILD function of HMMER and the PF00394, PF07731, PF07732, PF00255, PF13532, or PF00487 seed alignments of the Pfam database. The amino acid sequences of all hits with an e-value ≤ 0.01 were then collected for each HMM. In the case of laccases, the hits from the PF00394, PF07731, and PF07732 HMMs were pooled and dereplicated. Similarly, the hits from the PF04261, PF21105, and PF20628 HMMs were pooled and dereplicated as DyP-type peoxidases. Next, each set of proteins was queried against the entire Pfam v. 38.0 HMM database using the HMMSCAN function of HMMER, and the top scoring HMM for each protein was recorded. Proteins were then classified based on their top scoring HMM, after which the amino acid sequences of the proteins belonging to the five families of interest were collected. Proteins of interest were identified as those with a top hit of: PF00394, PF07731, or PF07732 for laccases; PF00255 for glutathione peroxidases; PF04261, PF21105, or PF20628 for DyP-type peroxidases; PF13532 for 2OG-Fe(II) oxygenases; and PF00487 for fatty acid desaturases.

**Text S3. Statistical analyses**

MAG abundance data (average sequencing depth), taxonomic classification of each MAG as determined by GTDB-Tk, and sample metadata files were imported to R and used to create a phyloseq object, after which the MAG abundance data was normalized per sample; the normalized data were used for all analyses except for the differential abundance analysis. To address questions related to the impact of insect species on microbiome composition, a phyloseq object containing the full dataset was prepared. In addition, the phyloseq object was subsequently subsetted by species (mealworms or superworms) to address questions related to the impact of diet on microbiome composition within an insect species. In addition, the GTDB-Tk data was modified to more closely match the NCBI taxonomy: (i) the class affiliation of the order *Burkholderiales* was changed to *Betaproteobacteria* from the classification of *Gammaproteobacteria* reported by GTDB-Tk; (ii) family affiliation of the genus *Brucella* was changed to *Bartonellaceae* from the classification of *Rhizobiaceae* reported by GTDB-Tk; (iii) and the order *Rhizobiales* was renamed to *Hyphomicrobiales*.

Shannon and Simpson diversity indices were estimated using the phyloseq::estimate_richness function. Statistical analysis of the log-transformed Shannon and Simpson diversity indices was performed using linear mixed models with the lmer function (lme4 package). In these models, fixed effects included insect host species, sequencing read count (millions of reads following QC), and their interaction, while insect cohort nested within species (six cohorts total, three for each species) was included as a random effect. For comparisons of diversity across diets within an insect species, diet and sequencing read count (millions of reads following QC) were included as fixed effects while insect cohort (three groups) was included as a random effect. Type II or Type III ANOVAs, as appropriate, were run using the Anova function of the car package, while estimated marginal means were calculated and pairwise comparisons performed using the emmeans function.

Bray-Curtis distances were calculated using the distance function of phyloseq from either MAG abundance data or after agglomerating the data at various taxonomic levels. The betadisper and permutest functions of the vegan package were used to test if the within-group dispersion per treatment was equal, while PERMANOVAs were run using the adonis2 function of vegan. Principle Component Analyses (PCoA) and capscale analyses were run using the ordinate function of phyloseq, and the results plotted using the plot_ordination function of phyloseq. For analyses across insect species, PERMANOVA was run with the distance matrix as the response variable and insect species, sequencing read count (millions of reads following QC), and their interaction as main effects. For analyses across diets within an insect species, PERMANOVA and capscale analyses were run with the distance matrix as the response variable and with diet, insect cohort (three groups), and sequencing read count as main effects.

Taxon relative abundance means and standard deviations were calculated with the dplyr package. Differential abundance of taxa across diets per insect species was examined using ANCOM-

BC2, either at the MAG level or after agglomerating the data at the species level. ANCOM-BC2 was run with a model that included diet as a fixed effect and insect cohort (three levels) as a random effect. P value adjustment was performed using Benjamini-Hochberg corrections. In addition, ANCOM-BC2 was run twice per dataset, once with the standard diet as the reference condition and once with the starvation diet as the reference condition. A taxon was considered differentially abundant if, in both comparisons, the adjusted p-value was ≤ 0.025 and the taxon passed the sensitivity analysis.
To evaluate the impact of the centrifugation step on the recovery of fungal CFUs, the ratio of bacterial to fungal CFUs was calculated, following which a paired, two-tailed t-test was run comparing the CFUs recovered with versus without centrifugation. For calculation of the bacteria to fungi ratios, values of zero were replaced with values of one, as this was the limit of detection.

**Text S4. Lower rate of mapping of superworm Illumina datasets to MAGs.**
The majority of reads from each metagenome sample mapped to the corresponding dereplicated set of MAGs, suggesting that the MAGs captured most of the bacterial community abundance and diversity in these environments. Consequently, we focused our analyses on the MAGs rather than performing analyses directly from the raw reads. However, the lower rate of read mapping for the superworm samples compared to the mealworm samples could suggest that by relying on the MAGs, we are missing a greater portion of the community diversity and abundance of the superworm samples compared to the mealworm samples. While we cannot rule this out, it could instead reflect that a lower percentage of insect host reads were removed during quality control for the superworm samples. The mealworm genome used for removing insect reads from the Illumina datasets was more complete than the superworm genome (99.1% versus 91.5% complete BUSCO endopterygota marker genes) [5]. As removal of contaminating host reads was performed through mapping of reads to the host genomes, the lower completeness of the superworm genome may mean that a higher percentage of host reads could have escaped detection and remained in the final read sets, which in turn, may explain why a lower fraction of reads mapped to the MAGs for this insect species.

**Text S5. Determination of bacterial and fungal colony forming units (CFUs)**
To enumerate bacterial and fungal CFUs from the gut microbiomes of mealworms and evaluate whether centrifugation of the collected microbiomes altered the relative recovery of bacteria versus fungi, two selective media were prepared. To select for bacteria, Tryptic Soy Agar (TSA; 30 g/L BD Bacto Tryptic Soy Broth catalog no. DF0370-07-5, 15 g/L agar) plates were prepared with 20 μg/mL of cycloheximide and 50 μg/mL of nystatin. To select for fungi, Potato Dextrose Agar (PDA; 24 g/L BD Difco Potato Dextrose Broth catalog no. DF0549179, 15 g/L agar) plates were prepared with 15 μg/mL of tetracycline, 15 μg/mL of nalidixic acid, and 25 μg/mL of chloramphenicol.

Insects were obtained and reared as described in the Materials and Methods subsection "Insect colonies and experimental design for metagenomic sequencing", except that the treatments were limited to only the standard diet and PS diet. After 10 days, the gut microbiomes of three replicates of 10 mealworms were collected in 1 mL of sterile 0.85% NaCl saline solution as described in the Materials and Methods subsection "Insect handling and DNA recovery". An aliquot of each sample was centrifuged at 1,000 × *g* for three minutes, after which the supernatant was transferred to a new tube. All samples (with or without centrifugation) were serial diluted in sterile saline and 100 μL of each dilution was spread plated onto both the TSA and PDA media described above. Plates were incubated at 28°C and colonies were counted daily for two and five days for the TSA and PDA plates, respectively.

We observed no effect of centrifugation on the ratio of bacterial versus fungal CFUs (paired 2-tailed t-test; p-value = 0.2), and we consistently recovered > 100,000 times more bacterial CFUs than fungal CFUs (**Table S1**). These results are consistent with the metagenomics data suggesting that the gut microbiota of mealworms and superworms are dominated by bacteria and have only a low abundance of fungal organisms. These results also indicate that centrifugation did not impact the relative recovery of bacteria versus fungi, and thus we kept the centrifugation step to reduce contamination of the cell suspensions with insect tissue.

**Table 1.** Colony forming units (CFUs) per mealworm digestive tract when plated on media selected for bacteria (TSA) or fungi (YPD).

| Diet | Centrifugation | Replicate | Bacterial CFU/insect | Fungal CFU/insect * | Ratio Bacteria / Fungi |
|---|---|---|---|---|---|
| Standard | No | 1 | 4,600,000 | 0 | 4,600,000 |
| Standard | No | 2 | 1,020,000 | 3 | 340,000 |
| Standard | No | 3 | 760,000 | 0 | 760,000 |
| Standard | Yes | 1 | 3,8000,00 | 0 | 3,800,000 |
| Standard | Yes | 2 | 250,000 | 0 | 250,000 |
| Standard | Yes | 3 | 220,000 | 0 | 220,000 |
| Polystyrene | No | 1 | 4,500,000 | 39 | 115,385 |
| Polystyrene † | No | 2 | 20,000 | 22 | 909 |
| Polystyrene | No | 3 | 760,000 | 3 | 253,333 |
| Polystyrene | Yes | 1 | 280,000 | 0 | 280,000 |
| Polystyrene | Yes | 2 | 340,000 | 1 | 340,000 |
| Polystyrene | Yes | 3 | 170,000 | 1 | 170,000 |

* Values of zero (0) were replaced with a value of one (1) during statistical testing, which was the limit of detection.

† T-tests were run both with (p-value = 0.38) and without (p-value = 0.20) this replicate included.

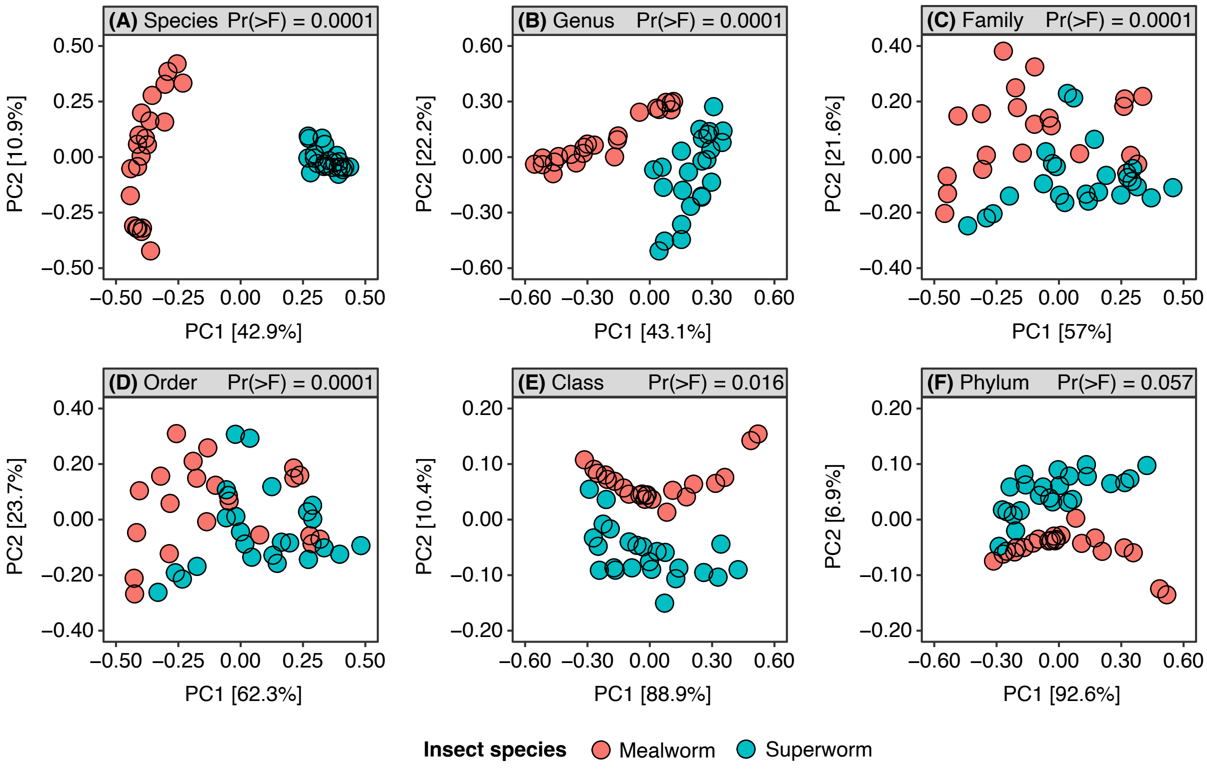


**Figure S1. Beta-diversity analyses of the mealworm and superworm gut bacterial communities at various taxonomic levels.** Principal Coordinate Analysis (PCoA) plots based upon the Bray-Curtis dissimilarity values between each sample are shown. Prior to the analyses, community abundances were summarized at the (**A**) species, (**B**) genus, (**C**) family, (**D**) order, (**E**) class, or (**F**) phylum level. Red dots represent communities from the mealworm samples while the blue-green dots represent communities from the superworm samples. The p-values were produced by running a PERMANOVA with the distance matrix as the response variable and insect species, read count, and their interaction as main effects using the adonis2 function of the vegan package in R. The percentages on the X and Y axes are the percent of the total variation explainable by principal component 1 (PC1) or 2 (PC2), respectively.

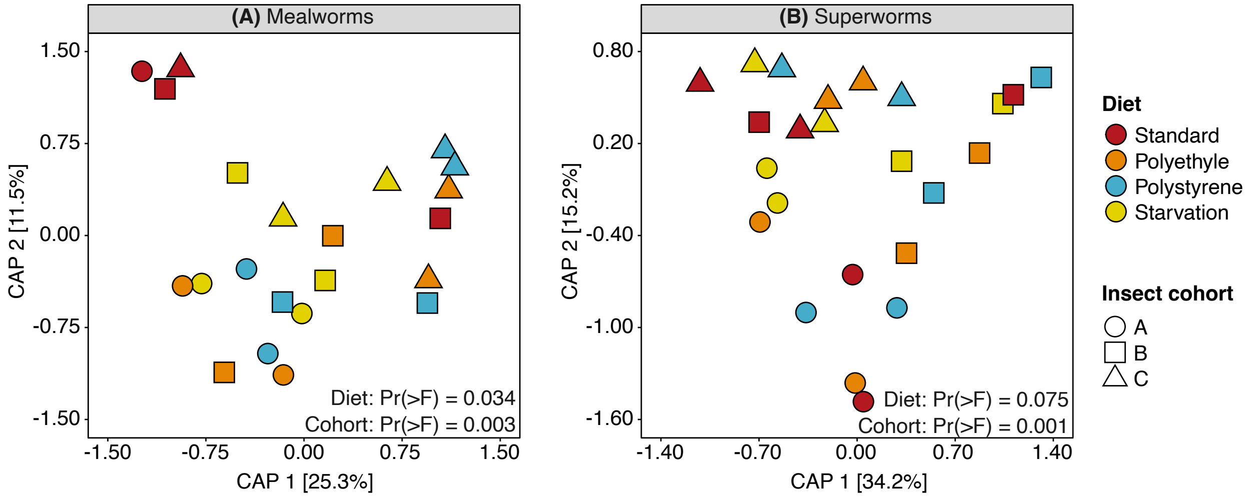


**Figure S2. Beta-diversity analyses of the mealworm and superworm gut bacterial communities across diets at the genus level.** Ordination plots produced from a capscale analysis run on Bray-Curtis distances are shown for (**A**) mealworms and (**B**) superworms. Bacterial community abundances were summarized at the genus level prior to running the capscale analyses. The p-values are from F-tests generated from running Type II ANOVAs on capscale analyses run with models that included diet, sequencing depth, and insect cohort as main effects.

### LEGENDS FOR DATASETS

**Dataset S1. Summary statistics for the metagenomic sequencing and assembly.** The column "% of raw read length" indicates the percentage of sequencing data that survived quality control. The column "% of reads mapping to bacterial MAGs" indicates the percentage of cleaned reads that mapped to the corresponding dereplicated MAG sets (i.e., the 53 mealworm MAGs or the 100 superworm MAGs). The column "% of full metagenome" indicates the percentage of the metagenomes that remained in the metagenomes when contigs < 2.5 kb in length were removed.

**Dataset S2. Summary statistics for the bacterial metagenome-assembled genomes (MAGs).** The "MAG name" refers to the full name used to label each MAG, while the "MAG ID" is a shorter identifier used in the manuscript. The "NCBI MAG ID" refers to the ID used while uploading the MAGs to NCBI; only MAGs with a completeness score > 90% and a contamination score < 5% were uploaded to NCBI. * The class affiliation of the order *Burkholderiales* was changed to *Betaproteobacteria* from the classification of *Gammaproteobacteria* reported by GTDB-Tk. Likewise, the family affiliation of the genus *Brucella* was changed to *Bartonellaceae* from the classification of *Rhizobiaceae* reported by GTDB-Tk, and the order *Rhizobiales* was renamed to *Hyphomicrobiales*.

**Dataset S3. Presence/absence of biosynthetic gene clusters (BGCs) in the 153 mealworm and superworm MAGs.** The first column gives the name and taxonomic classification of each MAG. The second column indicates the total number of BGCs producing distinct product types. The next five columns indicate the number of BGCs producing distinct product types within each BGC class. The following columns are named by BGC class followed by product type, and are binary columns indicating that a BGC encoding the given product type is present (1, green) or absent (0, yellow) from a given MAG.

**Dataset S4. Oxidative proteins identified in the secretome of the mealworm and superworm gut microbiomes**. The first column gives the internal protein name. The second column indicates which proteins belonged to a sequence similarity network (SSN) cluster that also included a protein previously implicated is PE or PS degradation, and for these proteins, provides the DOIs of the corresponding publications and the organism encoding the protein. The "BLAST results" section provides a summary of the results of performing a tBLASTn search of the 153 mealworm and superworm MAGs, indicating the MAG that was the top hit as well as the percent identity of the match and the query coverage. The "MAG taxonomic classification" section provides the taxonomic classification of the MAG that was the top hit in the tBLASTn search. The last column provides the amino acid sequence of the protein.
